# Generation and molecular characterization of CRISPR/Cas9-induced mutations in 63 immunity-associated genes in tomato reveals specificity and a range of gene modifications

**DOI:** 10.1101/835108

**Authors:** Ning Zhang, Holly M. Roberts, Joyce Van Eck, Gregory B. Martin

## Abstract

The CRISPR/Cas9 system is a powerful tool for targeted gene editing in many organisms including plants. However, most of the reported uses of CRISPR/Cas9 in plants have focused on modifying one or a few genes, and thus the overall specificity, types of mutations, and heritability of gene alterations remain unclear. Here we describe the molecular characterization of 361 T0 transgenic tomato plants that were generated using CRISPR/Cas9 to induce mutations in 63 immunity-associated genes. Among the T0 transformed plants, 245 carried mutations (68%), with 20% of those plants being homozygous for the mutation, 30% being heterozygous, 32% having two different mutations (biallelic) and 18% having multiple mutations (chimeric). The mutations were predominantly short insertions or deletions, with 87% of the affected sequences being smaller than 10 bp. The majority of 1 bp insertions were A (50%) or T (29%). The mutations from the T0 generation were stably transmitted to later generations, although new mutations were detected in some T1 plants. No mutations were detected in 18 potential off-target sites among 144 plants. Our study provides a broad and detailed view into the effectiveness of CRISPR/Cas9 for genome editing in an economically important plant species.

## Introduction

Derived from a native adaptive immune system in eubacteria and archaea, the CRISPR/Cas system enables the alteration of DNA sequences in many organisms to achieve precise gene modifications (Jaganathan et al., 2018). The most widely used *Streptococcus pyogenes* Cas9 (SpCas9) requires the 20-bp spacer sequence of a guide RNA (gRNA) to recognize a complementary target DNA site upstream of a protospacer adjacent motif (PAM) and generates a double-stranded breaks (DSB) near the target region (Xie and Yang, 2013). DSBs are repaired through either non-homologous end joining (NHEJ) or homology-directed recombination (HDR) resulting in small insertions/deletions (indels) or substitutions at the target region, respectively (Jinek et al., 2012). Compared to other genome editing tools such as zinc finger nucleases (ZFNs; Kim et al., 1996) and transcription activator-like effector nucleases (TALENs; Bogdanove and Voytas, 2011), CRISPR/Cas is more robust in that the Cas protein can theoretically bind to any genomic region preceding a PAM site and, importantly, target multiple sites simultaneously. However, the possibility of off-target mutations caused by CRISPR/Cas is a potential concern in both basic and applied research in plants, although it has been reported that off-target effects of CRISPR/Cas occur at a much lower frequency in plants than in mammals (Fu et al. 2013; Kuscu et al. 2014). The most effective way to minimize off-target mutations is to select a gRNA target with little or no homology to other genomic regions (Baltes and Voytas 2015). Other methods to reduce off-target mutations include using paired Cas9 nickases (Ran et al. 2013) or paired fusions of a catalytically dead Cas9 nuclease to the *Fok*I cleavage domain (Guilinger et al. 2014; Tsai et al. 2014).

As one of the most important vegetable crops in the world (Kimura and Sinha, 2008), tomato (*Solanum lycopersicum*) is an important source of health-promoting nutrients including vitamin C and E, minerals, and carotenes such as ß-carotene and lycopene (Canene-Adams et al., 2005). However, tomato production is threatened by many infectious diseases, including bacterial speck disease caused by *Pseudomonas syringae* pv. tomato (*Pst*), which can result in severe economic losses due to reduced yield and quality (Xin and He, 2013). A large number of candidate immunity-associated genes have been identified in tomato, but validation of the functional importance of these genes had been technically challenging before the emergence of CRISPR/Cas technology (Oh and Martin, 2011; Pedley and Martin, 2013; Pombo et al., 2014; Rosli et al., 2013). Although CRISPR/Cas has been used to modify genes with key roles in growth, development, and biotic and abiotic stresses in plants (Ito et al., 2015; Jacobs et al., 2017; Shimatani et al., 2017; Yu et al., 2017; Nekrasov et al., 2017; Rodriguez-Leal et al., 2017; D’Ambrosio et al., 2018; Li et al., 2018; Tashkandi et al., 2018; Hashimoto et al., 2018; Li et al., 2018; Zsogon et al., 2018; Ortigosa et al., 2019; Wang et al., 2019, Xu et al., 2019), all of the reported studies have focused on one or a few genes and could not provide broad insights into the specificity, types of mutations and heritability of genome editing by CRISPR/Cas9 in tomato.

Recently, we developed the Plant Genome Editing Database (PGED; http://plantcrispr.org/cgi-bin/crispr/index.cgi) which provides information about a collection of tomato lines with CRISPR/Cas9-induced mutations in immunity-associated genes (Zheng et al., 2019). In the present study, we molecularly characterized 361 T0 transgenic tomato plants that were generated using CRISPR/Cas9 to induce mutations in 63 candidate immunity-associated genes. To enhance the mutation efficiency and reduce the number of transformations needed, we evaluated gRNA efficiency by transient expression in tomato leaves and conducted tomato transformation with *Agrobacterium* pools containing 2-4 Cas9/gRNA constructs. This initial evaluation of gRNAs allowed us to select the most efficient ones for tomato stable transformation while using “*agrobacterium* pools” with various Cas9/gRNAs constructs shortened the time for generating multiple tomato mutant lines. We established an efficient CRISPR/Cas9 system to generate a large number of primary transgenic lines and we report for the first time a systematic investigation of the specificity of targeting, the types of mutations generated and the heritability of the mutations through multiple generations of tomato. Our CRISPR/Cas9-induced tomato mutant plants provide a powerful resource for better understanding the molecular mechanisms of plant-microbe interactions in the future.

## Materials and Methods

### Guide RNA (gRNA) design and construct development

All 20-nt gRNAs specific for the target genes were designed using the software Geneious R11 as described previously (Jacobs et al., 2017). The tomato (*Solanum lycopersicum*) reference genome sequence (SL2.5 or SL3.0) was used as an off-target database to score each gRNA (GN_19_ or gN_19_; “g” represents a manually added “G” to accommodate the transcription initiation requirement of the U6 promoter if the first nucleotide is not a G at target sites) preceding a PAM (NGG) sequence. For each gene, 2-3 gRNA targets with minimum off-target scores were designed. Single or multiple gRNA cassettes were cloned into a binary vector p201N:Cas9 by Gibson assembly as described previously (Jacobs and Martin, 2016). Colonies containing correct gRNA sequences were confirmed by PCR and Sanger sequencing.

### Evaluation of gRNA efficiency by agroinfiltration in tomato and *Nicotiana benthamiana* leaves

Each Cas9/gRNA vector was transformed into the *Agrobacterium tumefaciens* strain 1D1249 (Wroblewski et al., 2015) by electroporation. For agroinfiltration into tomato leaves, the bacterial cells containing different gRNA plasmids were grown in liquid YEP medium with 150 mg/L kanamycin overnight at 30°C. The bacterial pellet was collected and resuspended in an induction buffer containing 10 mM MgCl_2_, 10 mM MES (pH 5.7), and 200 μM acetosyringone (Sigma-Aldrich). Bacterial suspensions were adjusted to OD_600_ = 0.3 and incubated at room temperature for 2-5 h. The third and fourth leaves of 4-week-old tomato plants were infiltrated with needle-less syringes and the whole plant was then placed in a growth chamber with a temperature of 22-24°C, 16 h light/8 h dark photoperiod and 65% relative humidity. Three days later, a pool of six leaf discs were collected from three individual plants (two leaf discs from each of three plants) that had been infiltrated with the tested Cas9/gRNA vector, and used for genomic DNA extraction, PCR and sequencing. The web-based tool TIDE (https://tide.deskgen.com) was used to determine the mutation frequency induced by corresponding Cas9/gRNA vectors.

### Tomato transformation

Tomato transformation was performed either at the plant transformation facility at the Boyce Thompson Institute (BTI) or North Carolina State University(NCSU) (Gupta and Van Eck, 2016; Van Eck et al., 2019). Modifications of the transformation methods were made for Rio Grande (RG), including using 100 mg/l kanamycin for selection, and adding 0.1 mg/l indole-3-acetic acid (IAA) to the plant regeneration media (2Z, 1Z) and 1 mg/l IAA to the rooting medium. Each Cas9/gRNA vector was first electrotransformed into *Agrobacterium tumefaciens* LBA4404 (BTI), AGL1 (BTI) or GV3101(pMP90) (NCSU). In most cases, 2-4 *Agrobacterium* culture preparations (of the same *Agrobacterium* strain), each carrying a different Cas9/gRNA construct, were pooled together and used for transformation to minimize the number of experiments.

Tomato genotypes RG-PtoR or RG-prf3 were used for transformation if not specifically labeled (**Table 1**).

**Table 1.** Mutation rates and mutation types in T0 transgenic plants. See also Table S3.

| Target genes | Solyc # | # of transgenic plants | # of edited plants | Mutation rate (%)<sup>a</sup> | Mutation types<sup>b</sup> |
| --- | --- | --- | --- | --- | --- |
| ADE | Solyc05g005700 | 10 | 5 | 50 | 3 homo; 1 hetero; 1 chimeric |
| AOX | Solyc08g075550 | 9 | 8 | 89 | 2 homo; 2 biallelic; 3 hetero; 2 chimeric |
| APE | Solyc11g018800 | 1 | 1 | 100 | 1 biallelic |
| Aquaporin 1/2/3<sup>c</sup> | Solyc11g069430 (Aquaporin 1) | 12 | 10 | 83 | 5 homo; 2 biallelic; 10 hetero; 2 chimeric |
|  | Solyc06g074820 (Aquaporin 2) |  |  |  |  |
|  | Solyc08g066840 (Aquaporin 3) |  |  |  |  |
| BHLH | Solyc03g114230 | 3 | 3 | 100 | 3 biallelic |
| BSK830 | Solyc12g099830 | 8 | 6 | 75 | 2 biallelic; 3 hetero; 1 chimeric |
| BSK830 (Hawaii 7981)<sup>d</sup> | Solyc12g099830 | 6 | 5 | 83 | 3 biallelic; 1 hetero; 1 chimeric |
| Bti9-interactor | Solyc09g008010 | 1 | 1 | 100 | 1 homo |
| Bti9ab | Solyc07g049180 | 4 | 3 | 75 | 2 homo; 1 biallelic |
| CathepsinB1 | Solyc02g076980 | 29 | 20 | 69 | 8 homo; 8 biallelic; 2 hetero; 2 chimeric |
| CathepsinB2 | Solyc02g077040 | 11 | 8 | 73 | 2 homo; 3 biallelic; 3 hetero |
| CORE | Solyc03g096190 | 7 | 6 | 86 | 1 homo; 4 biallelic; 1 hetero |
| Drm-3 | Solyc01g099840 | 3 | 3 | 100 | 2 biallelic; 1 hetero |
| EDS1 | Solyc06g071280 | 2 | 2 | 100 | 1 homo; 1 chimeric |
| ERF5 | Solyc03g093560 | 7 | 5 | 71 | 2 biallelic; 1 hetero; 2 chimeric |
| Fen | Solyc05g013290 | 18 | 13 | 72 | 5 biallelic; 7 hetero; 1 chimeric |
| Fen (RG-pto11)<sup>e</sup> | Solyc05g013290 | 15 | 9 | 60 | 2 homo; 4 biallelic; 5 chimeric |
| Fls2.1 | Solyc02g070890 | 4 | 4 | 100 | 3 homo; 1 hetero |
| Fls3 | Solyc04g009640 | 11 | 7 | 64 | 5 homo; 2 hetero |
| LRRXII-1 | Solyc06g076910 | 5 | 1 | 20 | 1 biallelic |
| LRRXII-2 | Solyc04g012100 | 4 | 3 | 75 | 1 biallelic; 2 hetero |
| Mai1 | Solyc04g082260 | 4 | 1 | 25 | 1 hetero |
| Mai5 | Solyc10g085990 | 11 | 8 | 73 | 2 homo; 5 hetero; 1 chimeric |
| MAP3Ka | Solyc11g006000 | 6 | 6 | 100 | 4 biallelic; 2 hetero |
| Min7 | Solyc12g017830 | 5 | 4 | 80 | 1 biallelic; 1 hetero; 2 chimeric |
| MKK1 | Solyc12g009020 | 2 | 2 | 100 | 1 biallelic; 1 chimeric |
| MKK2 | Solyc03g123800 | 6 | 5 | 83 | 1 homo; 4 biallelic |
| MKK4 | Solyc03g097920 | 2 | 1 | 50 | 1 hetero |
| MKKK15 | Solyc02g065110 | 8 | 6 | 75 | 4 hetero; 2 chimeric |
| MKKK66 | Solyc08g081210 | 3 | 1 | 33 | 1 hetero |
| MLO16 | Solyc11g069220 | 4 | 4 | 100 | 1 biallelic; 2 hetero; 1 chimeric |
| NOD | Solyc11g008200 | 8 | 7 | 88 | 1 homo; 4 hetero; 2 chimeric |
| NPR1 | Solyc07g040690 | 5 | 1 | 20 | 1 hetero |
| NRC1/2/3<sup>f</sup> | Solyc01g090430 (NRC1) | 8 | 5 | 63 | 1 homo; 3 biallelic; 1 hetero |
|  | Solyc10g047320 (NRC2) |  |  |  |  |
|  | Solyc05g009630 (NRC3) |  |  |  |  |
| PAD4 | Solyc02g032850 | 7 | 1 | 14 | 1 biallelic |
| PBCP | Solyc03g116690 | 3 | 2 | 67 | 1 homo; 1 biallelic |
| PBL-T1 | Solyc09g007170 | 5 | 3 | 60 | 1 biallelic; 2 chimeric |
| PBL-T2 | Solyc01g067400 | 2 | 2 | 100 | 2 hetero |
| Peptide Transporter 3 | Solyc05g009500 | 4 | 3 | 75 | 1 biallelic; 1 hetero; 1 chimeric |
| Permease Transporter | Solyc03g005820 | 4 | 2 | 50 | 1 homo; 1 biallelic |
| PGA1<sup>f</sup> | Solyc05g005560 | 5 | 2 | 40 | 2 hetero |
|  | Solyc05g005570 |  |  |  |  |
| Pic1 | Solyc07g066260 | 5 | 4 | 80 | 1 biallelic; 2 hetero; 1 chimeric |
| PR1b | Solyc00g174340 | 6 | 4 | 67 | 3 biallelic; 1 chimeric |
| Propep1 | Solyc04g072310 | 2 | 2 | 100 | 2 chimeric |
| RALF1 | Solyc01g067900 | 3 | 3 | 100 | 1 homo; 2 biallelic |
| RALF2 | Solyc01g099520 | 10 | 7 | 70 | 1 homo; 3 hetero; 3 chimeric |
| RbohB | Solyc03g117980 | 8 | 3 | 38 | 1 homo; 2 biallelic |
| SAG101-1/2<sup>f</sup> | Solyc02g069400 (SAG101-1) | 3 | 2 | 67 | 1 homo; 1 biallelic |
|  | Solyc02g067660 (SAG101-2) |  |  |  |  |
| SIMIo1/9<sup>f</sup> | Solyc04g049090 (SIMIo1) | 10 | 9 | 90 | 1 homo; 2 biallelic; 3 hetero; 6 chimeric |
|  | Solyc06g010030 (SIMIo9) |  |  |  |  |
| SOBIR/SOBIR-like<sup>f</sup> | Solyc06g071810 (SOBIR) | 16 | 9 | 56 | 2 homo; 1 biallelic; 4 hetero; 3 chimeric |
|  | Solyc03g111800 (SOBIR-like) |  |  |  |  |
| Solute Transporter 2 | Solyc05g005950 | 4 | 2 | 50 | 1 biallelic; 1 hetero |
| STP13 | Solyc09g075820 | 2 | 1 | 50 | 1 homo |
| TFT1 | Solyc11g010470 | 3 | 3 | 100 | 1 biallelic; 2 hetero |
| TFT10 | Solyc04g076060 | 4 | 2 | 50 | 2 chimeric |
| TFT7 | Solyc04g074230 | 2 | 2 | 100 | 2 hetero |
| Wak1 | Solyc09g014720 | 5 | 5 | 100 | 2 homo; 2 biallelic; 1 chimeric |
| WRKY11 | Solyc08g006320 | 2 | 2 | 100 | 2 chimeric |
| WRKY9b | Solyc08g067360 | 4 | 1 | 25 | 1 biallelic |

<sup>a</sup>Mutation rate (%) = number of plants with mutations/number of total transgenic plants. Mutations were analyzed by Geneious R11 and TIDE.
<sup>b</sup>Homo: homozygous mutation; hetero: heterozygous mutation; <sup>c</sup>Three gRNA cassettes were cloned into one p201N:Cas9 plasmid; <sup>d</sup>Tomato genotype Hawaii 7981 was used for transformation; <sup>e</sup>Tomato genotype RG-pto11 was used for transformation; <sup>f</sup>Specific gRNA for each gene was independently cloned into the p201:Cas9 vector and the two or three gRNA/Cas9 constructs were pooled together for tomato transformation.

### Genotyping and mutation analysis

Genomic DNA was extracted from tomato cotyledons or young leaves using a modified CTAB method (Murray and Thompson, 1980). The existence of T-DNA was confirmed by PCR using primers listed in **Table S6**. To determine the mutation specificity, genomic regions flanking the target site of each gene were amplified with specific primers (see PGED; http://plantcrispr.org/) and sequenced by Sanger sequencing. TIDE was used to rapidly evaluate the mutated allelic sequences using the sequencing files (.ab1 format), especially for PCR amplicons of biallelic, heterozygous, or chimeric mutants whose mutation length and frequency cannot be determined due to superimposed sequencing chromatograms.

### Off-target evaluation

To evaluate potential off-target mutations caused by gRNAs in CRISPR-induced mutant plants, twelve gRNAs were selected and used as queries to search for potential off-target sites across the tomato genome with up to 4-nt mismatches and 1-nt indel by the software Geneious R11 or with up to 3-nt mismatches by a web tool Cas-OFFinder. Each off-target site was given a score based on how similar it was to the protospacer of gRNAs. A higher score for an off-target site indicated a higher similarity to the original target site and a higher likelihood to cause off-target mutations. A shortlist of potential off-target sites of each gRNA queried was generated by selecting their relatively high-scoring off-target sites predicted by either Geneious R11 or Cas-OFFinder (**Table 3**). Similar to mutation genotyping described above, genomic regions flanking the putative off-target sites were amplified with specific primers (**Table S7**) and PCR amplicons were sequenced to detect if off-target mutations were induced in those regions.

## Results

### CRISPR/Cas9 targeting of immunity-associated genes in tomato

To study the efficiency and specificity of genome editing in tomato by CRISPR/Cas9 and to better understand plant-pathogen interactions, we generated a collection of tomato lines with targeted CRISPR/Cas9-induced mutations in genes that have been implicated in the immune response. Candidate genes were selected based on previous studies involving RNA-Seq, biochemical approaches, virus-induced gene silencing (VIGS) or yeast two-hybrid (Y2H) screens (Rosli et al., 2013; Pombo et al., 2014; Zeng et al., 2012; Giska and Martin, 2019); orthologs of immunity-associated genes reported in other plant species such as Arabidopsis and rice were also included (Hutin et al., 2015; Xin et al., 2016; Shimizu et al., 2010; Stegmann et al., 2017; Yamada et al., 2016).

For each candidate gene, at least two gRNAs targeting different DNA sites were designed and separately cloned into a Cas9-expressing binary vector p201N:Cas9 (Jacobs and Martin, 2016). The gRNAs were designed to be highly specific at target sites and their predicted off-target sites contained at least one nucleotide mismatch in the seed sequence (the last 12 nucleotides preceding the PAM) or two nucleotide mismatches against the full 20-bp protospacer (although some gRNAs were designed to intentionally modify multiple homologs simultaneously). Most Cas9/gRNA constructs in this study had only one gRNA expression cassette per plasmid, except one construct that contained three gRNA cassettes targeting three *Aquaporin transporter* genes (**Table 1**).

### Evaluation of gRNA effectiveness by agroinfiltration in tomato and *Nicotiana benthamiana* leaves

To initially evaluate the effectiveness of gRNAs and subsequently enhance the mutation rate in stably transformed tomato plants, 195 gRNAs were tested for their ability to cause mutations using *Agrobacterium* infiltration (agroinfiltration) in tomato leaves (**Fig. 1A**; **Table S1**). After agroinfiltration, DNA was isolated from the leaf tissue and the genomic region spanning each target site was PCR amplified, sequenced, and analyzed with a web-based tool called Tracking of Indels by Decomposition (TIDE; https://tide.deskgen.com; Brinkman et al., 2014) to calculate the mutation frequency. gRNAs with mutation frequency >0 (p<0.0001) were considered to be effective in inducing mutations in this assay while those with mutation rate = 0 were considered ineffective. A total of 61.5% of the tested gRNAs were effective in inducing transient mutations in tomato leaves (**Fig. 1B**). Among these, 96% had a mutation rate greater than 0% but less than 10% in this assay, while only five gRNAs (4%) had a mutation frequency over 10% (**Fig. 1C**).

**Figure 1.**
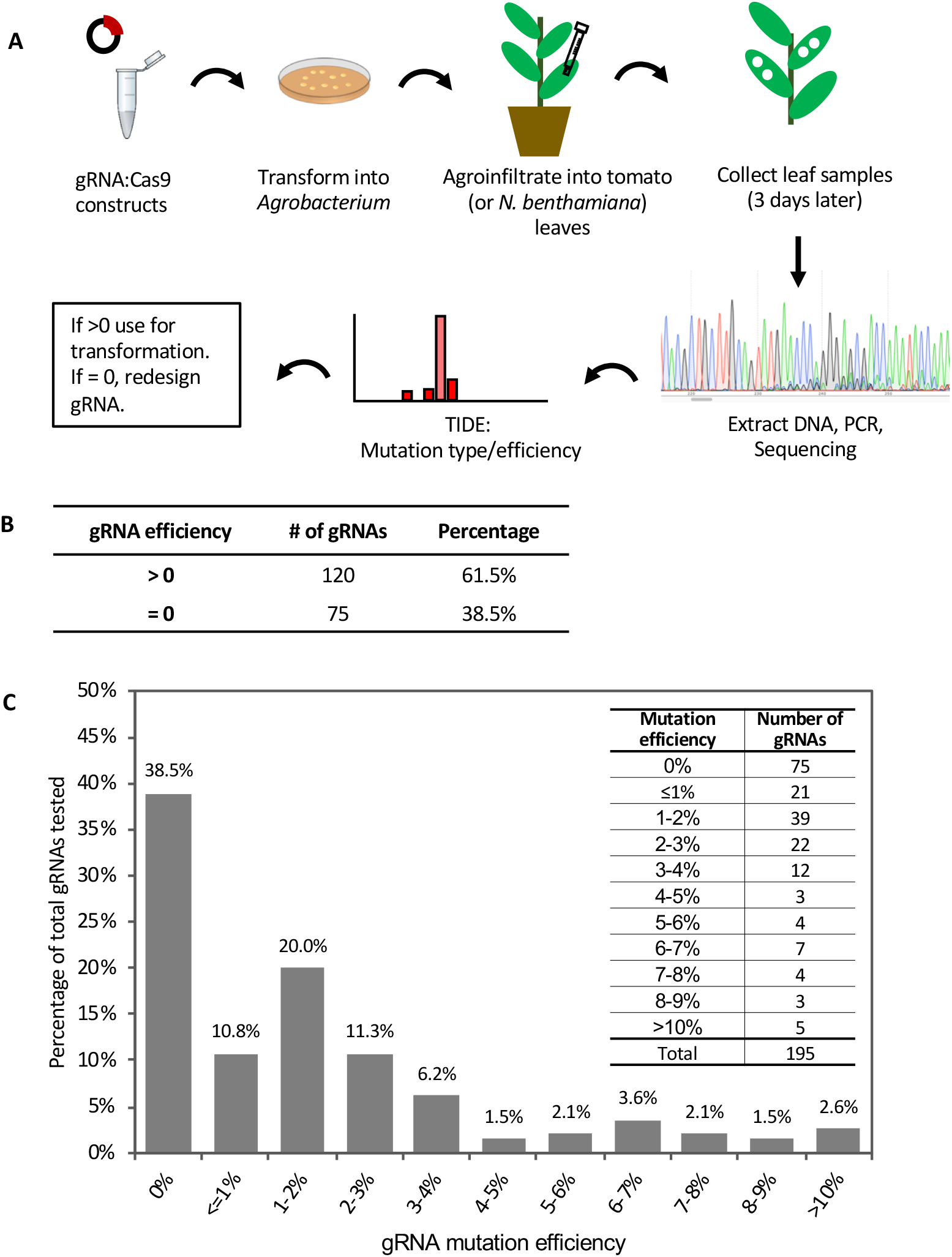
Evaluation of gRNA-mediated mutation efficiency by agroinfiltration in tomato leaves. See also Figure S1 and Table S1. (**A**) Schematic showing the workflow of gRNA evaluation by agroinfiltration. **(B)** Summary of gRNA efficiency tested by agroinfiltration. **(C)** The distribution of mutation efficiencies of the 195 gRNAs. Inset on the top right shows the number of gRNAs in each mutation efficiency range. TIDE (https://tide.deskgen.com) was used to calculate mutation efficiency by identifying the predominant types of insertions and deletions (indels) in the DNA of a targeted cell pool.

Agroinfiltration of tomato leaves is not very efficient and to test whether this affected our estimate of gRNA mutation efficacy, we evaluated the mutation efficiency of two of the gRNAs (Bti9ab-1 and Drm3-1) which each have identical target sites in both tomato and *Nicotiana benthamiana* leaves (**Fig. S1**). The mutation frequency induced by these two gRNAs in tomato was much lower than in *N. benthamiana* (**Fig. S1A**). In addition to these two gRNAs, we also tested another four gRNAs that each target two of the four homologs of the *Mai1* gene in *N. benthamiana* (Roberts et al., 2019; **Fig. S1B**). In *N. benthamiana*, the majority of the gRNA targets showed a mutation frequency of 10-40%, while a small number had a mutation frequency less than 5% (**Fig. S1B-C**). These observations suggest that the inefficiency of agroinfiltration in tomato leaves probably leads to an underestimate of the true efficacy of gRNAs for generating mutations. This is supported by later observations in which some very low-efficient gRNAs were very effective in inducing mutations in stably-transformed tomato plants (**Table S2**).

In most cases, we selected the most efficient gRNAs for subsequent stable transformation in tomato, however, some low-efficiency gRNAs were also used if limited gRNAs could be designed for a particular target gene. Most gRNAs that were effective in inducing mutations in the agroinfiltration transient assay also induced mutations in stable transgenic seedlings, with one exception where a gRNA that had a 4.4% mutation frequency in the transient assay did not edit target genes in two stably-transformed plants (**Table S2**). It was not possible, however, to directly compare gRNA efficiency in the transient assay with that in stable transformation, as other factors such as the bias of gRNA transformation into plants using ‘*Agrobacterium* pools’ and the total number of regenerated transgenic seedlings varied from gene to gene in stable transformation.

### CRISPR/Cas9-induced mutations in T0 transgenic plants

A total of 361 putative primary (T0) transgenic tomato plants were regenerated by *Agrobacterium tumefaciens*-mediated stable transformation. To confirm the mutated sequence(s) in each plant, genomic regions spanning the target sites were PCR amplified and sequenced. All five possible genotypes, that is, wild-type, homozygous for the mutation, biallelic (a different mutation in each allele), heterozygous for the mutation, or multiple mutations (chimeric), were detected in our stably transformed tomato plants (**Table 1**). Direct sequencing of PCR amplicons containing biallelic, heterozygous, or chimeric mutations resulted in superimposed sequencing chromatograms, which made it difficult to determine specific mutation types and mutation frequency in those plants. To resolve this problem, TIDE was used to rapidly determine the mutated allelic sequences using the sequencing file (.ab1 format) with superimposed chromatograms (Brinkman et al., 2014), thus avoiding tedious and expensive cloning and multi-clone sequencing for mutation analysis.

Of the 361 T0 plants, 245 were found to have modifications at the target site(s) within 63 genes (**Table 1**; **Table S3**). Most of the lines had only one CRISPR-induced mutation in one gene per plant, while a few had mutations in two or even three genes (the latter cases occurred when using Agrobacterium pooling – see below). We identified only one mutant event for some of the targeted genes while for others up to 20 independent mutant events were generated (**Table 1**). Overall, the average editing efficiency (the number of edited plants / the number of transgenic plants) by CRISPR/Cas9 in tomato in our experiments was 68% (**Fig. 2A**), although the mutation rate varied over a wide range from 14% to 100% from target gene to target gene in different mutant lines (**Table 1**). All four mutation types (homozygous, biallelic, heterozygous, or chimeric) were observed in several mutant lines that had sufficient independent mutant events, while there was a bias of mutation types in some mutant lines, probably due to the limited number of transgenic events generated (**Table 1**).

**Figure 2.**
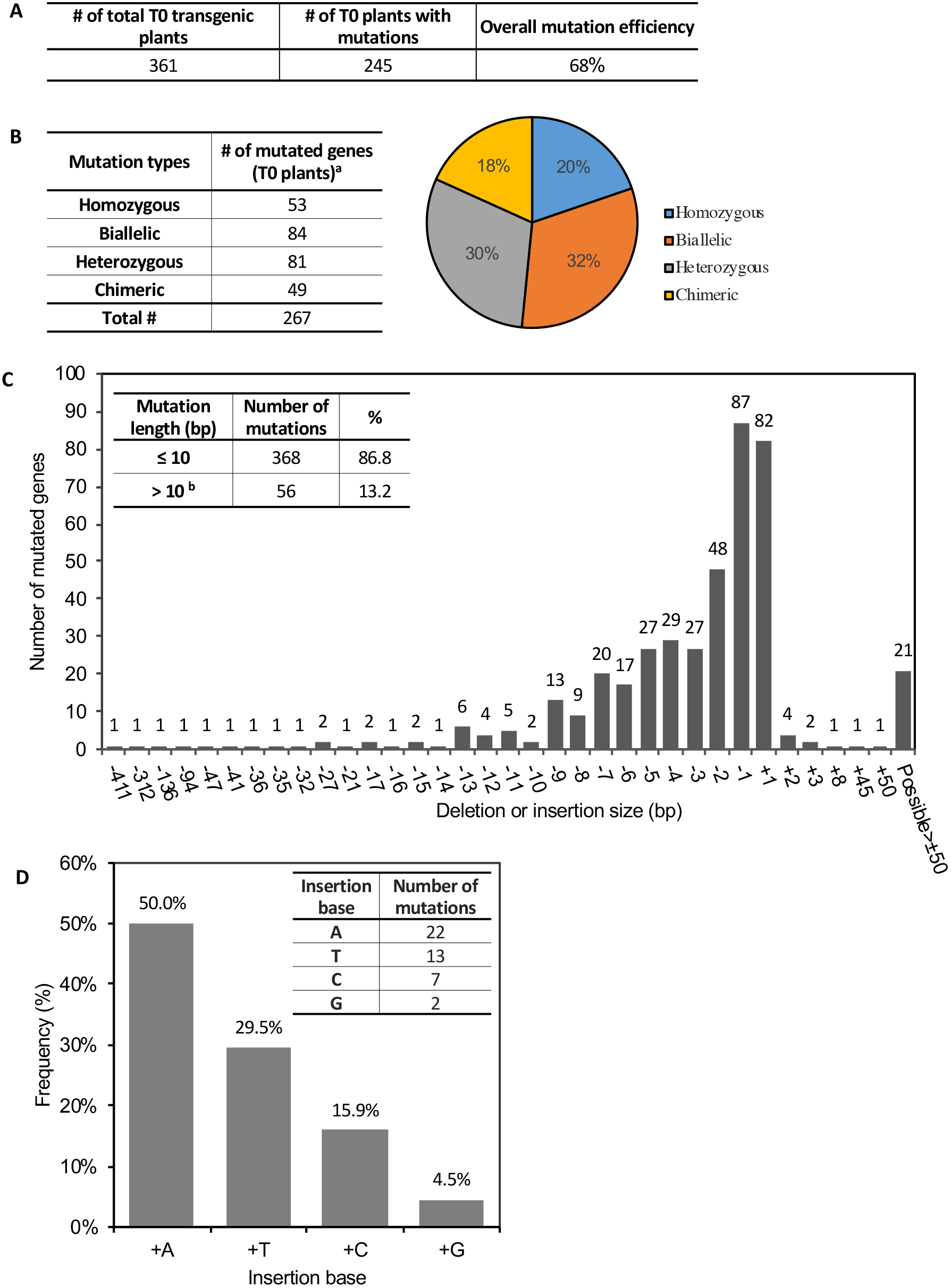
CRISPR/Cas9-induced gene mutations in T0 transgenic plants. See also Table S3. (**A**) The average mutation rate induced by CRISPR/Cas9 in T0 plants. **(B)** Summary of CRISPR-induced mutation types and their frequency in T0 plants. Left: Number of genes modified with the corresponding mutation type; Right: Percentage of genes harboring the corresponding mutation type. ^**a**^ Some plants have multiple target genes in one plant. **(C)** Frequency of each insertion or deletion mutation. x-axis: number of base pairs (bp) deleted (−) or inserted (+) into target sites. Inset on the left top shows the percentage of mutations ≤10 bp or >10 bp. ^**b**^ All “possible > ± 50 bp” in the figure are included in >10 bp. TIDE only calculates mutation length ≤50 bp. **(D)** Percentage of different bases in the 1-bp insertion mutations. Inset at the right top shows the number of mutations with each type of inserted base.

We analyzed the distribution of the four mutation types in all the 245 T0 plants at the 267 mutated target sites (some plants had more than one target site) and found that the percentage of homozygous, biallelic, heterozygous, or chimeric mutation was 20%, 32%, 30%, and 18%, respectively (**Fig. 2B**). In particular, plants having homozygous or biallelic mutations accounted for 52% of the total. These mutants and their progenies can be used directly for phenotype screening because no wild-type alleles are present, thus speeding the research process by saving time for further genotyping in the next generation. The most common mutation alterations induced by CRISPR/Cas9 were deletions or insertions, with 87% of these modifications at the target sites being less than 10 bp (**Fig. 2C**). The proportion of deletion mutations was 77%, and the deletion length spanned a wide range from 1 bp to over 400 bp. Of all the mutations, the most abundant modification was 1-bp deletion or insertion (**Fig. 2C**). For these, A- and T-insertions accounted for 79.5%, while G-insertions accounted for only 4.5% (**Fig. 2D).** Base substitutions in combination with indels were also detected, but at a much lower frequency. Only three independent mutant events (two were the same mutation type) harbored a nucleotide substitution in one copy of the target genes at the positions 5-bp preceding the PAM, along with a short insertion or deletion at the target site (**Fig. S2**).

Multiplex editing of three *Aquaporin transporter* (*AquaT*) genes by using one Cas9/gRNA construct was also tested. Three individual promoter-gRNA expression cassettes (in the order *Aqua1-Aqua2-Aqua3*) were assembled into the p201N:Cas9 vector (**Fig. S3A**) as previously reported (Jacobs et al., 2017). Ten of the 12 regenerated transgenic plants were edited, including three single mutants, four double mutants, and three triple mutants (having mutations in all three genes). Interestingly, all the three single mutants knocked out *AquaT1*, while three double mutants modified *AquaT1* and *AquaT2* and one edited *AquaT1* and *AquaT3* simultaneously. Three plants had mutations in all the three *AquaT* genes together (**Fig. S3B**). These data suggested the position of the gRNA cassette in the vector may affect its mutation rate, considering that all gRNAs were efficient enough to induce mutations once transformed into plants.

### Heritability of the mutations

To evaluate the heritability of mutations in T0 plants, a large number of T1 and some T2 plants were generated and examined for their genotypes at the target sites. Most of the same genotypes from T0 plants were transmitted to plants in later generations, although we did not record segregation ratios in the progenies. Of note, no new mutations or reversions to wild-type were found in the progeny of any homozygous T0 plants, indicating all the homozygous mutations occurred in the transformed embryogenic cells before the first division. However, we did observe novel genotypes in a small percentage of T1 or T2 plants whose progenitor (T0 plants) harbored biallelic, heterozygous or chimeric mutations (**Table 2**). In particular, a homozygous mutation (−265 bp) was detected in the progeny of a *NRC2* primary transgenic plant with a “+1 bp/+2 bp” biallelic mutation. It was unlikely that the new −265 bp modification at the target site was created by further modification of the existing +1 bp or +2 bp mutations in the progenitor, as the insertions occurred at the fourth nucleotide upstream of the PAM, where a 1-bp mismatch in the seed sequence of the gRNA protospacer can severely impair or completely abrogate the Cas9/gRNA functionality (Jiang and Doudna, 2017). One possible explanation for this observation is that the biallelic T0 plant (+1 bp/+2 bp) was a chimera and the new mutation may derive from chimeric tissue of the T0 plant (Zhang et al., 2014).

**Table 2.** New genotypes detected in T1 or T2 plants.

| T0 mutation type<sup>a</sup> | Generation | Plants | Gene Solyc# | Mutation<sup>b</sup> | Is T-DNA present? |
| --- | --- | --- | --- | --- | --- |
| **Biallelic** | **T0** | **NRC2-E4** | **Solyc10g047320** | **+1 bp/+2 bp** | **Yes** |
|  | T1 | NRC2-E4-P2 | Solyc10g047320 | +1 bp/large deletion<sup>c</sup> | No |
|  | T2 | NRC2-E4-P2-2 | Solyc10g047320 | -265 bp/-265 bp | No |
|  | T2 | NRC2-E4-P2-4 | Solyc10g047320 | +1 bp/+1bp | No |
| **Heterozygous** | **T0** | **Mai1-E10** | **Solyc04g082260** | **-4 bp/WT** | **Yes** |
|  | T1 | Mai1-E10-P20 | Solyc04g082260 | +1 bp/WT | Yes |
| **Heterozygous** | **T0** | **CathepsinB2-E8** | **Solyc02g077040** | **-4 bp/WT** | **Yes** |
|  | T1 | CathepsinB2-E8-P2 | Solyc02g077040 | -2 bp/-2 bp | No |
|  | T1 | CathepsinB2-E8-P10 | Solyc02g077040 | -1 bp/WT | No |
| **Heterozygous** | **T0** | **Min7-E6** | **Solyc12g017830** | **+1 bp/WT** | **Yes** |
|  | T1 | Min7-E6-P4 | Solyc12g017830 | +1 bp/-3 bp | Yes |
|  | T1 | Min7-E6-P8 | Solyc12g017830 | -1 bp/WT | No |
| **Heterozygous** | **T0** | **NRC2-E1** | **Solyc10g047320** | **-3 bp/WT** | **Yes** |
|  | T1 | NRC2-E1-P6 | Solyc10g047320 | -3 bp/-5 bp | Yes |
|  | T1 | NRC2-E1-P15 | Solyc10g047320 | -3 bp (58%); -6 bp (32%) +1 bp (3.6%) | No |
| **Chimeric** | **T0** | **MKKK15-E2** | **Solyc02g065110** | **-1 bp (10.4%); +1 bp (2.1%); WT (85%)** | **Yes** |
|  | T1 | MKKK15-E2-P2 | Solyc02g065110 | -5 bp(60%); -1 bp (31%); +1 bp (4.7%) | Yes |
|  | T1 | MKKK15-E2-P3 | Solyc02g065110 | -5 bp (20%); WT (74%) | No |
|  | T1 | MKKK15-E2-P7 | Solyc02g065110 | +1 bp (77.6%); -5 bp (4.7%); WT (9.5%) | Yes |
| **Chimeric** | **T0** | **MKK1-E1** | **Solyc12g009020** | **-7 bp (8.1%); -2 bp (33.3%); WT (48.5%)** | **Yes** |
|  | T1 | MKK1-E1-P3 | Solyc12g009020 | -2 bp/-4 bp | No |
| **Chimeric** | **T0** | **Min7-E5** | **Solyc12g017830** | **-1 bp (35.9%); +1 bp (9%); WT (49.4%)** | **Yes** |
|  | T1 | Min7-E5-P1 | Solyc12g017830 | -1 bp/-3 bp | Yes |
|  | T1 | Min7-E5-P2 | Solyc12g017830 | -6 bp (64.5%); +1 bp (19%); -1 bp (8.8%);<br>-7 bp (3.4%) | Yes |
|  | T1 | Min7-E5-P3 | Solyc12g017830 | -5 bp/-6 bp | Yes |
|  | T1 | Min7-E5-P7 | Solyc12g017830 | -9 bp (71.3%); +1 bp (18.2%); -1 bp (3.6%) | Yes |
| **Chimeric** | **T0** | **ADE-E2** | **Solyc05g005700** | **-4 bp (20%); -5 bp (40.4%); -13 bp (13.9%)** | **Yes** |
|  | T1 | ADE-E2-P1 | Solyc05g005700 | WT/WT | No |
|  | T1 | ADE-E2-P20 | Solyc05g005700 | -4 bp/-9 bp | No |

<sup>a</sup>For each mutant event, only the plants harboring different genotypes from their T0 progenitors are listed. Unlisted T1 or T2 progeny have the same genotypes as those in T0 plants.
<sup>b</sup>Number of base pairs (bp) deleted (-) or inserted (+) into target sites; WT, wild-type. If no percentage is shown, the two genotypes are around 50%:50%
<sup>c</sup>TIDE only detects indels ≤50 bp.

We also observed new genotypes in the progeny of some T0 plants harboring heterozygous or chimeric mutations (**Table 2**) even though most of the progeny still possessed the same genotypes as the progenitor line. In some of these cases, CRISPR/Cas9 continued to modify the wild-type allele of the target gene in the progeny if the parent plants still contained a wild-type allele and the Cas9/gRNA expression cassette (**Table 2**). In other cases, unexpected genotypes were detected in some mutant lines including Mai1-E10, CathepsinB2-E8, Min7-E6, and ADE-E2 (**Table 2**). For instance, although the ADE-E2 T0 plant was chimeric (−4 bp/−5 bp/-13 bp) without a wild-type allele, we identified one T1 plant that was azygous (two copies of wild-type allele) and another one with a novel biallelic mutation (−4/−9 bp). Another example is the CathepsinB2-E8 T0 plant which had a heterozygous (−4 bp/WT) mutation. However, −2 bp homozygous and −1 bp/WT heterozygous mutations were detected in later generations. The unexpected new genotypes discussed above revealed that the one leaf/cotyledon sample may not reveal all the genotypes in the whole plant if the mutant is chimeric. Therefore, for T0 edited plants without any wild-type allele, it will still be useful to perform genotyping in subsequent generations to obtain homozygous mutants without the presence of Cas9/gRNA.

### Specificity of CRISPR-induced gene modifications in tomato

Mutations in unintended sequences (off-target mutations) is a possible concern in both functional genomics studies and plant breeding. To evaluate potential off-target effects by CRISPR/Cas9 in our tomato lines, we first evaluated the specificity of 12 gRNAs of Cas9 by Geneious R11 (https://www.geneious.com; Kearse et al., 2012) and Cas-OFFinder (Bae et al., 2014). The putative off-target sites predicted by Cas-OFFinder were then manually checked using JBrowse (https://solgenomics.net/jbrowse_solgenomics/) to confirm their locations in the tomato genome. The presence of a PAM was required for the site to be considered a candidate site. These gRNAs were selected for off-target analysis because morphological defects were observed in one or more mutant lines induced by these gRNAs (**Fig. S4**; **Table S4**). A total of 18 possible off-target sites of the 12 gRNAs were identified and off-target mutations were examined in 12 T0 plants, 68 T1 plants and 44 T2 plants by PCR and Sanger sequencing (**Table 3**). No off-target modifications were discovered in the tested plants with or without Cas9, indicating our gRNAs and CRISPR-mediated mutations are highly specific.

**Table 3.**
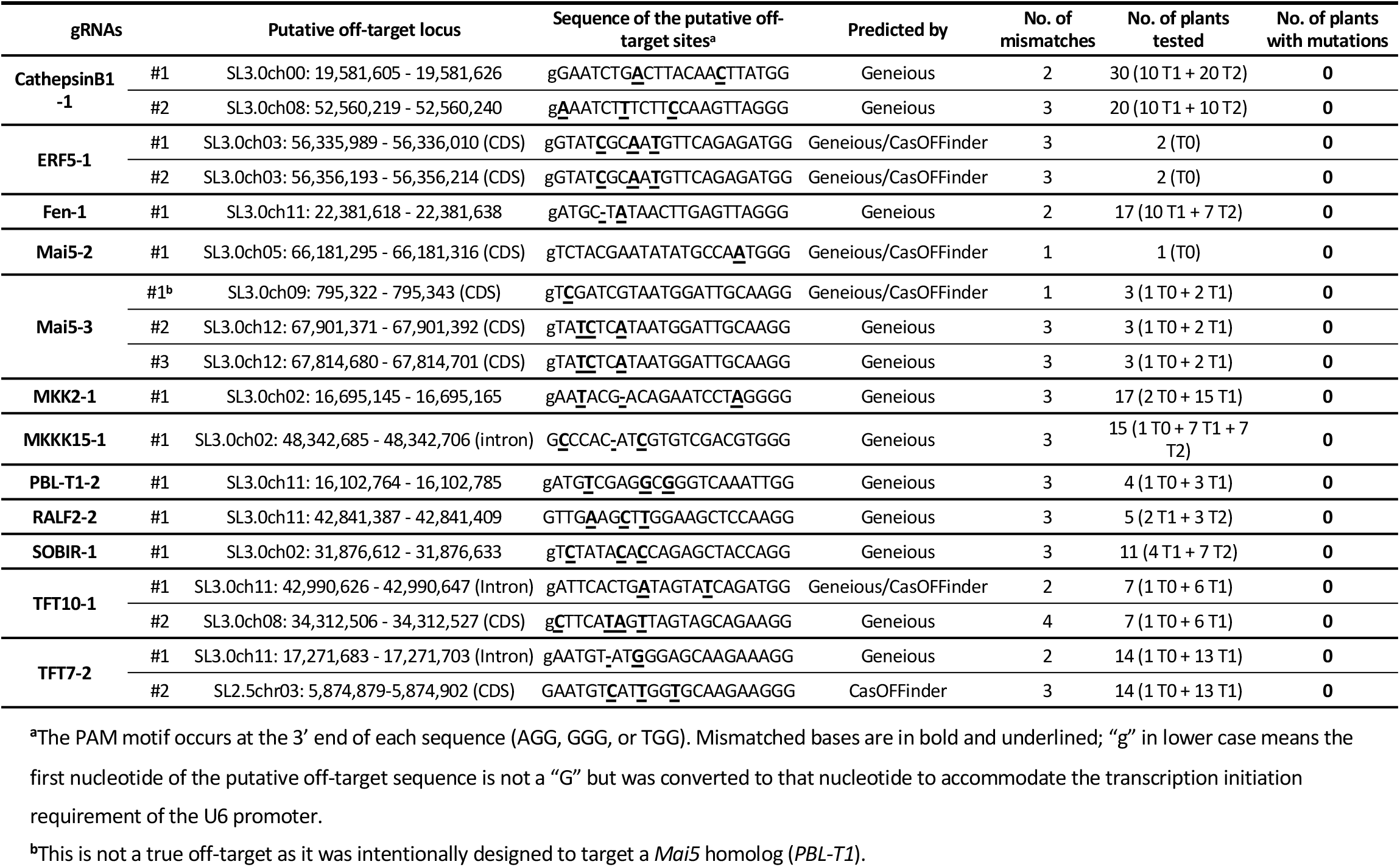
Examination of possible off-target mutations caused by 12 selected gRNAs in multiple generations. See also Table S4 and Table S7.

Another way to evaluate the specificity of CRISPR/Cas9 is to test the efficiency of gRNAs with a few mismatches against the target sequence in the protospacer. One of our gRNAs targeted two tomato homologs, *Mai5* and *PBL-T1* (bothMai5/PBL-T1; 5’-**g**TAGATCGTAATGGATTGCA-3’; the first nucleotide “C” was converted to “G” to accommodate the transcription initiation requirement of the U6 promoter). The designed 20-bp protospacer sequence exactly matched the target site in *Mai5* but had one mismatch in *PBL-T1* at the third nucleotide from the 5’ end (5’-gT<u>**C**</u>GATCGTAATGGATTGCA-3’). We generated five T0 plants that contained the bothMai5/PBL-T1 gRNA construct, all of which had edits in *Mai5* but not in *PBL-T1*, indicating the one mismatch (along with the first nucleotide at the 5’ end) in *PBL-T1* appeared to significantly affect Cas9 binding and cleavage activity at the target site. Another gRNA, targeting the tomato *FLS2.2* gene, did not induce targeted modifications in any of the 10 transgenic plants, possibly due to a 1-bp mismatch in the seed region of the gRNA in the Heinz 1706 reference genome (GTCATCAACAT<u>**C**</u>TCGCTTGT) as compared to Rio Grande-PtoR (GTCATCAACAT<u>**T**</u>TCGCTTGT). The reference genome was used for gRNA design and RG-PtoR was used for tomato transformation as it contains the resistance gene *Pto* for investigating *Pto*-mediated immunity in our mutants. This further indicates that CRISPR/Cas9 is highly specific, with even one mismatch in the gRNA rendering the site inaccessible to the Cas9/gRNA complex.

### Tomato transformation with *Agrobacterium* pools

Tomato transformation is a lengthy and labor-intensive process. In an approach to minimize the number of transformation experiments needed, three to four *Agrobacterium* culture preparations each carrying a different gRNA construct were pooled and used for a single transformation experiment (**Fig. 3A**). Of the 79 T0 plants generated, 58 plants (73%) contained precise modifications in one or more of the target genes. In terms of the number of target sites edited by CRISPR/Cas9 with pooled gRNAs, 48 (82.8%) of the T0 plants had mutations in just one gene, while 9 plants (15.5%) had mutations in two and 1 (1.7%) had mutations in three genes (**Fig. 3A**). Among these T0 plants, 83.5% contained one gRNA cassette and 15.2% contained two different gRNA cassettes, while no plants recovered contained more than two gRNA cassettes integrated into the genome (**Fig. 3B**). Interestingly, one mutant plant did not show detectable integration of the T-DNA sequence (expressing *Cas9* and gRNA) but had a mutated gene, suggesting that transient expression of the Cas9/gRNA occurred in this plant. Additionally, we found another type of transient mutation in 8 T0 plants at 10 different target sites (**Table S5**). In these plants, a Cas9/gRNA expression cassette was integrated into the plant genome, as confirmed by PCR and Sanger sequencing, but the gRNA detected was not the one that induced the mutation in the plant (**Table S5**), suggesting that the mutation was caused by another transiently expressed Cas9/gRNA.

**Figure 3.**
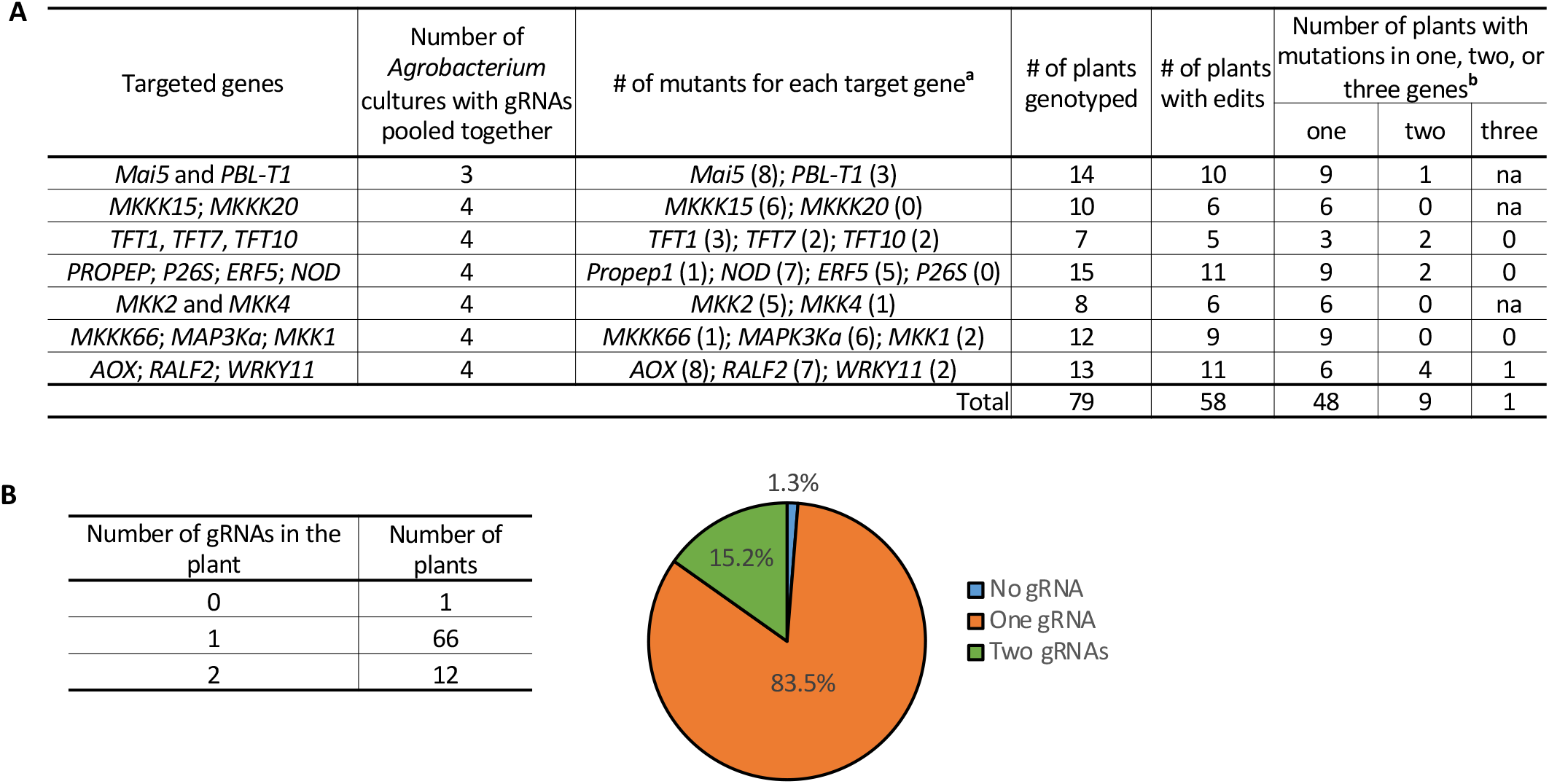
Tomato transformation with ‘Agrobacterium pools’. See also Table S5. (**A**) An example of a tomato transformation experiment designed to target 2-4 genes by using 3-4 pooled *Agrobacterium* cultures with each culture carrying a different gRNA. ^**a**^ Number of plants with mutations in the target gene is shown in parentheses; some plants had mutations in multiple genes. ^**b**^ Number of genes modified in the plants. “na” not applicable, since less than 3 genes were targeted in the experiment. **(B)** Left: Number of gRNAs detected in a single mutant plant by PCR and Sanger sequencing. Right: Percentage of T0 plants harboring no, one or two gRNAs.

## Discussion

Our effort to generate a large number of CRISPR/Cas9-induced tomato mutants targeting immunity-associated genes demonstrates that this mutation approach is efficient and robust for gene editing in tomato. Importantly, gene modifications mostly occurred in germline cells and were stably inherited in subsequent generations, similar to those in rice (Zhang et al., 2014) but not as in Arabidopsis, in which most mutations in T0 plants were somatically modified if a strong constitutive promoter was used to regulate the Cas9 expression (Feng et al., 2014; Feng et al., 2018). However, we detected a greater range of deletion and insertion lengths than observed in rice (Zhang et al., 2014), in which only 1-bp insertions and fewer deletion lengths were found, possibly due to different intrinsic DNA repair mechanisms between these two species. These differences could also be due to other factors including different transformation methods or culture conditions, and different sets of target genes that tolerate different degrees of mutations.

Base substitutions induced by the CRISPR/Cas9 system in tomato were very rare in our study. We frequently observed single nucleotide polymorphisms (SNPs) between Rio Grande (used for transformation) and the tomato reference genome (Heinz 1706), but these SNPs were due to natural variation, not mutagenesis, as confirmed by sequencing of the gene regions from untransformed plants. Most of these SNPs were located outside of the protospacer sequence of the gRNAs, and to date we have only found one gRNA (targeting *FLS2.2*) which had a mismatch in the seed region that inhibited the Cas9 binding and cleavage at the target site. Therefore, it is reasonable to use the tomato reference genome as the template for gRNA design and subsequent mutation genotyping in transgenic Rio Grande and likely other tomato cultivars.

Various morphological phenotypes were detected in some mutants compared to wild-type plants. Some of these abnormal phenotypes were associated with all the mutation events occurring in a specific gene, strongly supporting that the mutation itself is responsible for the altered plant growth or development. However, some mutant lines showed unusual morphology associated with certain mutation events but not all, possibly indicating that another off-target mutation occurred in these plants. We therefore investigated a large number of these plants but did not find any evidence of off-target mutations, suggesting other mutations, if they exist, were either caused by tissue culture or *Agrobacterium* transformation, or spontaneous mutations during seed propagation (Tang et al., 2018). Our observations are consistent with previous reports that CRISPR/Cas9 causes few off-target mutations in plants including Arabidopsis (Feng et al., 2014), rice (Zhang et al., 2014; Tang et al., 2018), tomato (Nekrasov et al., 2017), cotton (Li et al., 2019), and maize (Young et al., 2019). True off-targets reported previously in plants showed high sequence homology to the original protospacer sequence of gRNAs (Tang et al., 2018), which can be easily avoided by designing highly specific gRNAs using tools such as Geneious and Cas-OFFinder. Based on our data we devised a rule to avoid off-target effects of CRISPR/Cas9 by designing gRNAs whose highest scored potential off-target sites have at least a 1-nt mismatch in the seed sequence or 2-nt mismatches in the full protospacer sequence.

Surprisingly, we found new mutations in the progeny of some T0 plants that did not contain any wild-type allele. These new mutations did not appear to be derived from existing mutations in the T0 plants, as the Cas9-induced modifications were located within the seed sequence of the gRNA protospacer and as little as 1-bp mismatch in the seed sequence can dramatically impair the Cas9 binding and cleavage activity (Jinek et al., 2012). Therefore, we believe the new mutations were derived from chimeric tissue from the T0 plant that was not detected with the one cotyledon/leaf sample we used for mutation genotyping. We are currently advancing lines that have biallelic or heterozygous mutations, or that were chimeric to develop homozygous plants without the presence of Cas9/gRNA sequence. These plants will be used to investigate whether the mutations affect the plant immune response, especially to *P. syringae* pv. tomato.

## Acknowledgments

We thank Brian Bell, Jay Miller and Nick Vail for plant care, Patricia Keen, Michelle Tjahjadi, and Sergei Krasnyanski for tomato transformation, and Colleen Conger and Elizabeth Trost for assistance with mutation genotyping. The authors declare no conflicts of interest. Funding was provided by National Science Foundation grants IOS-1732253 (JVE) and IOS-1546625 (GBM).

## Author contributions

GBM and NZ conceived and designed the experiments. NZ designed gRNAs, constructed vectors, performed genotyping in T0 plants and other generations, and analyzed the data. HMR performed genotyping of mutants in some T1 and T2 plants. JVE optimized the plant transformation protocol and guided the transformation experiments. NZ and GBM interpreted the data and wrote the manuscript.

## Conflict of interest

The authors declare that they have no conflict of interest.

## Supplemental Information

**Figure S1.**
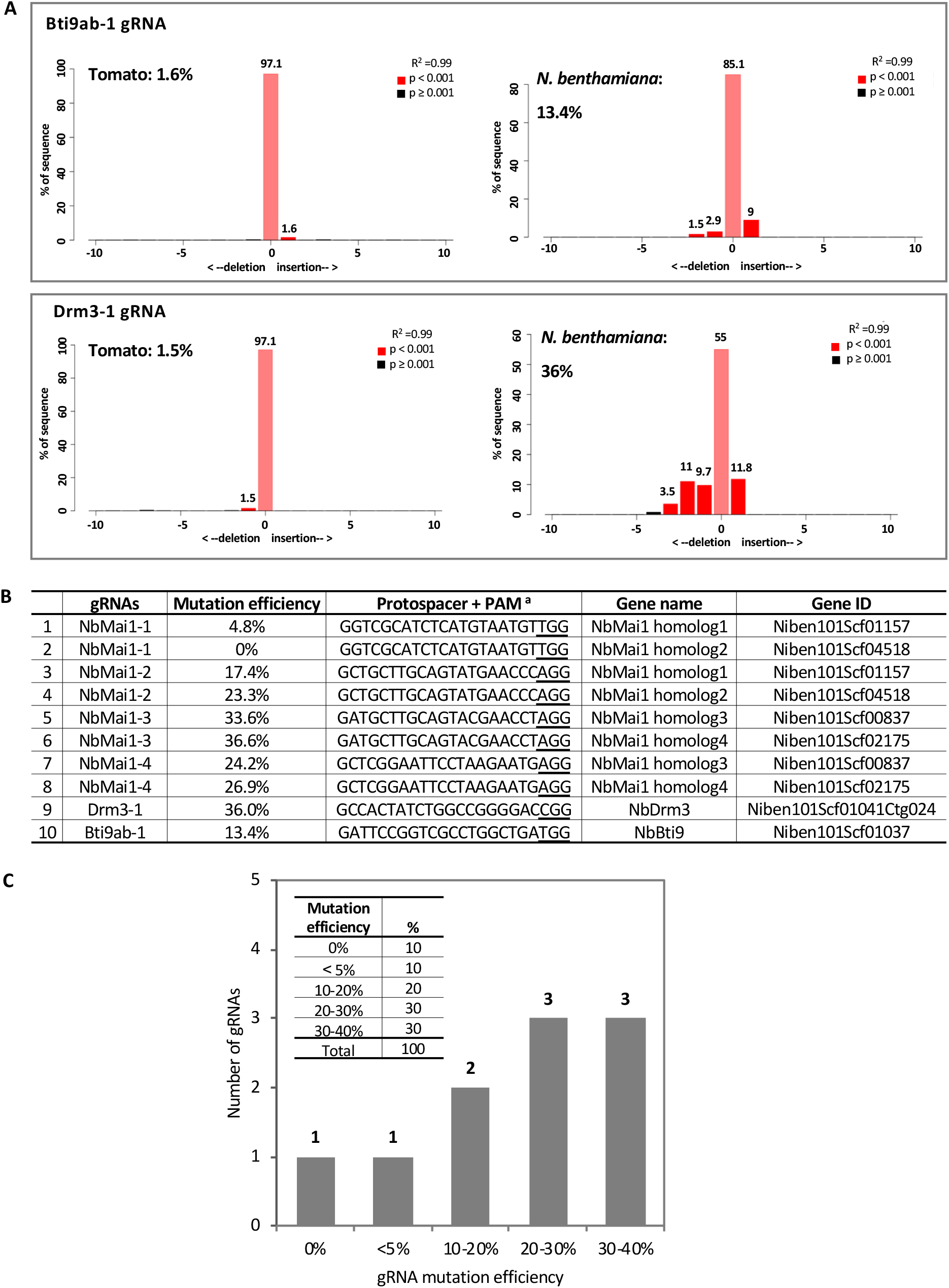
Evaluation of gRNA efficiency by agroinfiltration in *Nicotiana benthamiana* leaves. Related to Figure 1. (**A**) Comparison of the efficiency of the same gRNA to create mutations in tomato or *N. benthamiana*. Agroinfiltration and mutation analysis were the same as described in Fig. 1. **(B)** Efficiency of 10 gRNAs in *N. benthamiana* by agroinfiltration. ^**a**^ PAM (NGG) are underlined. **(C)** Distribution of gRNA mutation frequency in *N. benthamiana*. Inset on the top left shows the percentage of each mutation frequency range.

**Figure S2.**
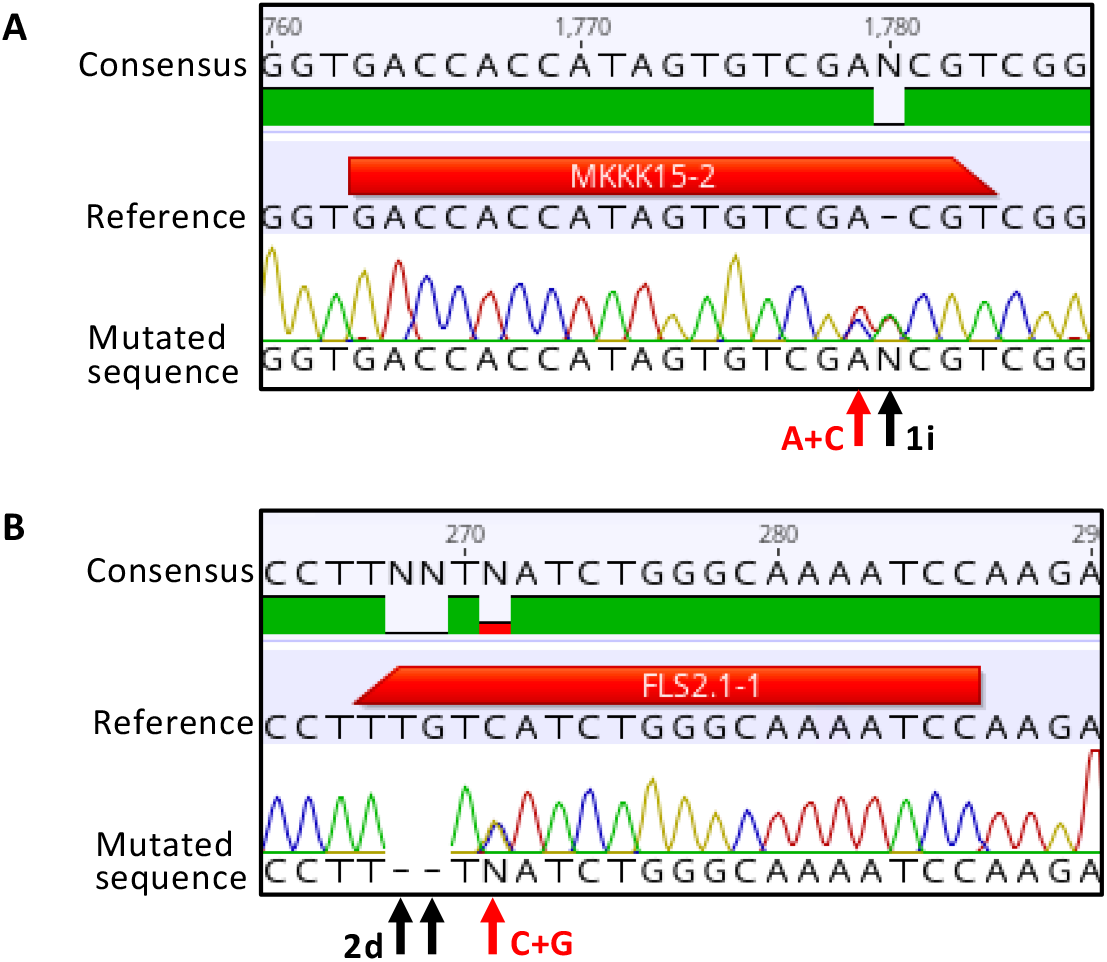
Base substitutions in tomato mutant plants by CRISPR/Cas9. (**A**) Mutation caused by the MKKK15-2 gRNA was 1-bp insertion plus a “A to C” substitution in one copy of the allele. 1i: 1-bp insertion. (**B**) Mutation caused by the FLS2.1-1 gRNA was 2-bp deletion plus a “C to G” substitution in one copy of the allele. 2d: 2-bp deletion.

**Figure S3.**
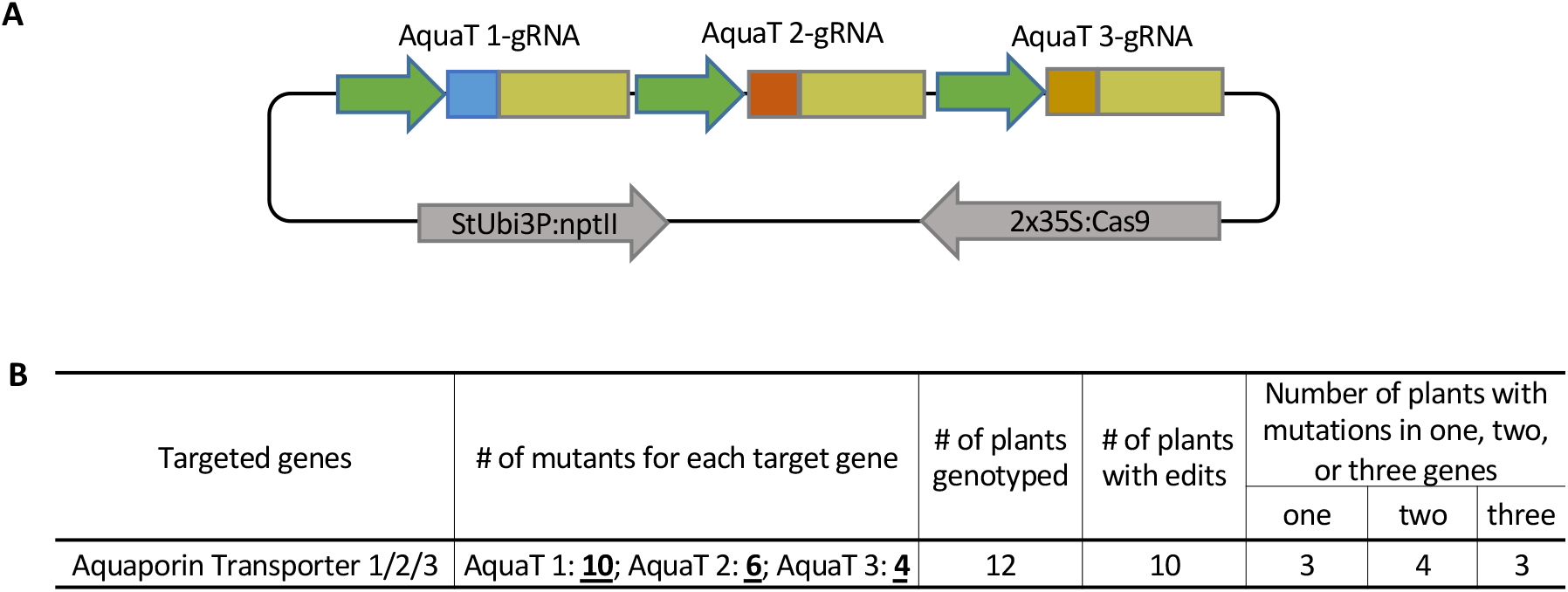
Multiplex genome editing by CRISPR/Cas9 in tomato. (**A**) Schematic shows the order of three gRNAs cassettes targeting *Aquaporin Transporter* genes in the p201N:Cas9 vector. AquaT: Aquaporin Transporter. **(B)** Summary of multiplex editing of three AquaT genes in tomato.

**Figure S4.**
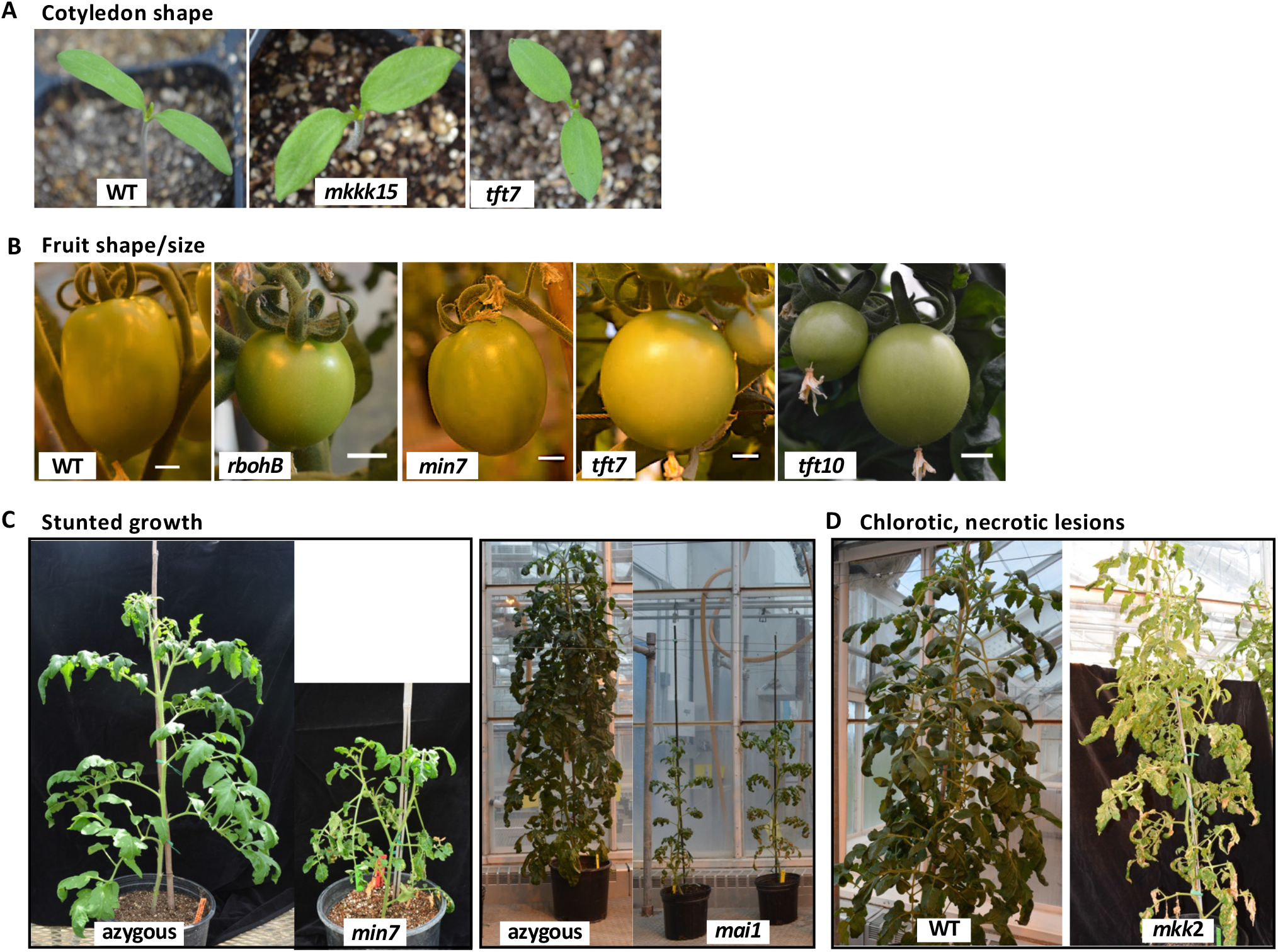
Morphology defects in some CRISPR-induced tomato mutants knocking out immunity-associated genes. (**A**) Changed cotyledon shape from elongate (wild-type, WT) to ovate (*mkkk15/tft7* mutants). Cotyledon size is similar between WT and mutant plants. (**B**) Change in fruit shape and size in some of the mutant plants. Scale bar: 1 cm. (**C**) Some mutants show stunted growth. Azygous plants are from the same transformation event but contain two copies of the wild-type alleles. Azygous and mutant plants are the same age. (**D**) The mkk2 mutant showed chlorotic and necrotic lesions in leaves.

**Table S4.**
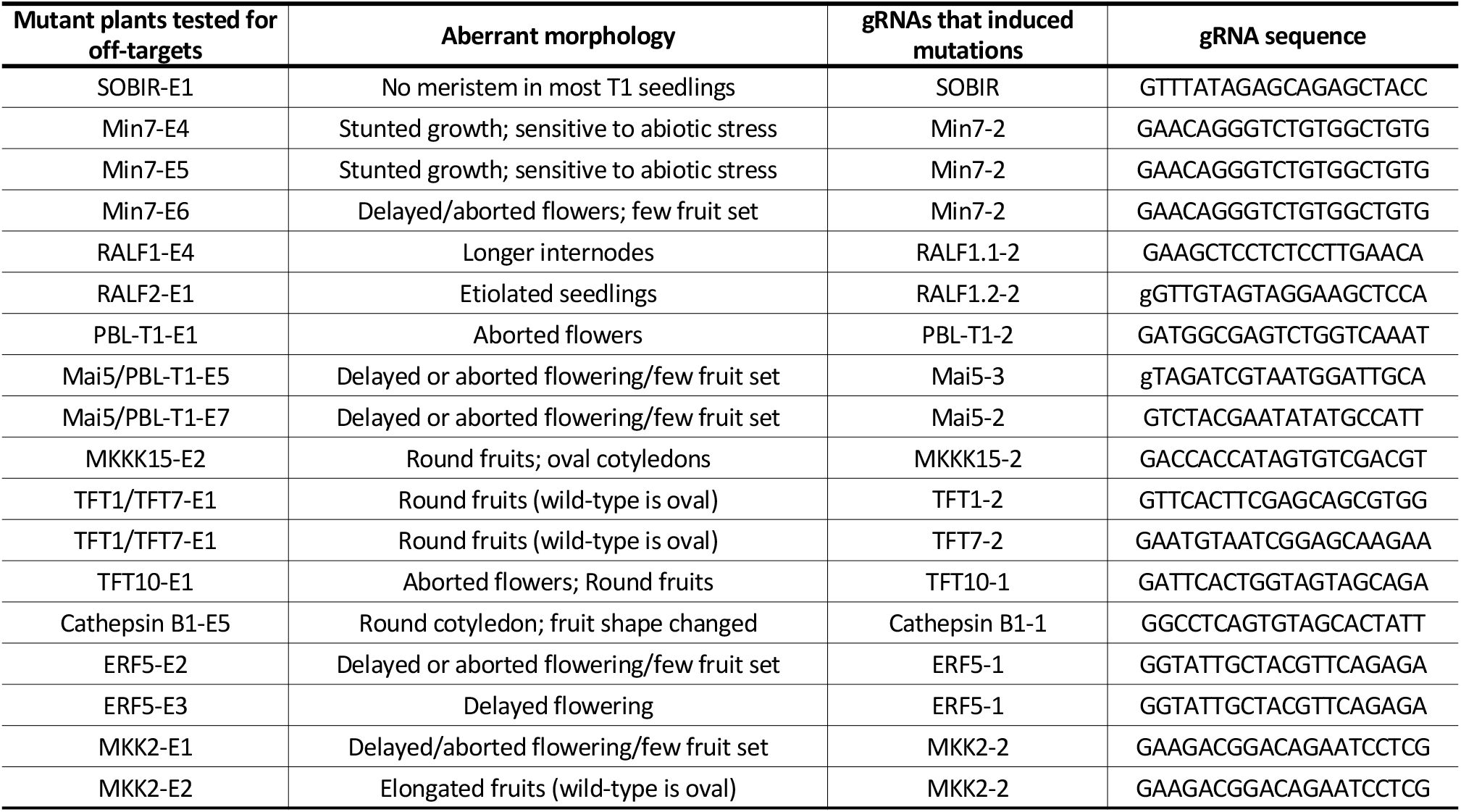
Tomato mutant events selected for off-target analysis showed morphology defects. Related to Table 3.

**Table S5.** Transient genome editing by CRISPR/Cas9 in tomato stable transformation. Related to Figure 3.

| Plants | Target gene | Gene Solyc# | Mutations | gRNAs that induced mutations | gRNAs detected in the plants<sup>a</sup> |
| --- | --- | --- | --- | --- | --- |
| 1 | AOX | Solyc08g075550 | -1 bp/WT | AOX-2 | WRKY11-4 |
|  | AOX | Solyc08g075550 | +1 bp/WT | AOX-1 | WRKY11-4 |
|  | RALF2 | Solyc01g099520 | -2 bp/WT | RALF1.2-1 | WRKY11-4 |
| 2 | ERF5 | Solyc03g093560 | +1 bp/-11 bp/WT | ERF5-1 | WRKY11-4 |
| 3 | MAP3Ka | Solyc11g006000 | +1 bp/+50 bp | MAP3Ka-3 | MKKK66-1 and MKK1-1 |
| 4 | MKKK15 | Solyc02g065110 | -5 bp/WT | MKKK15-2 | MKKK20-1 |
| 5 | MKKK15 | Solyc02g065110 | -5 bp/WT | MKKK15-2 | MKKK20-1 |
| 6 | MKKK15 | Solyc02g065110 | +1 bp/WT | MKKK15-2 | PCR failed |
| 7 | PBL-T1 | Solyc09g007170 | +1 bp/-5 bp | PBL-T1-1 | Both Mai5/PBL-T1 |
| 8 | RALF2 | Solyc01g099520 | +1 bp/WT | RALF1.2-1 | AOX-1 |

<sup>a</sup> These gRNAs were pooled together with other gRNAs for one transformation experiment, detected in transgenic plants by PCR and sequencing, but did not induce mutations at the target site.

**Table S6.**
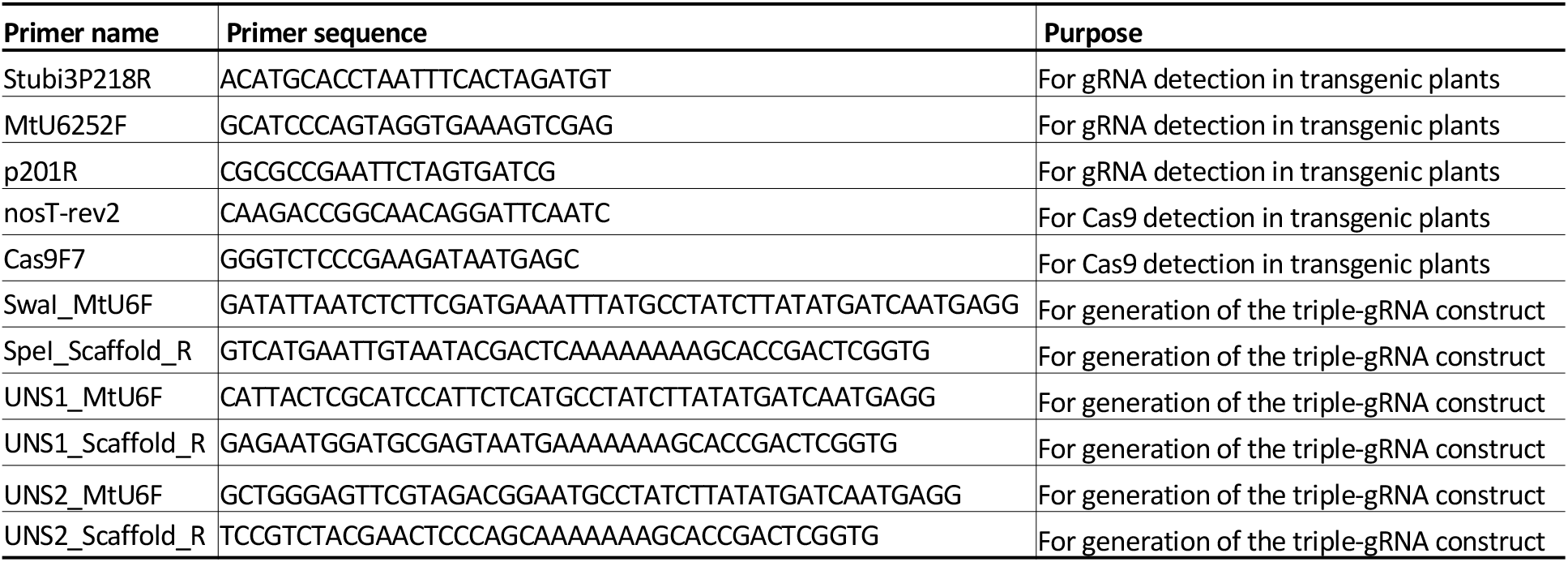
Primers used for cloning and genotyping.

**Table S7.**
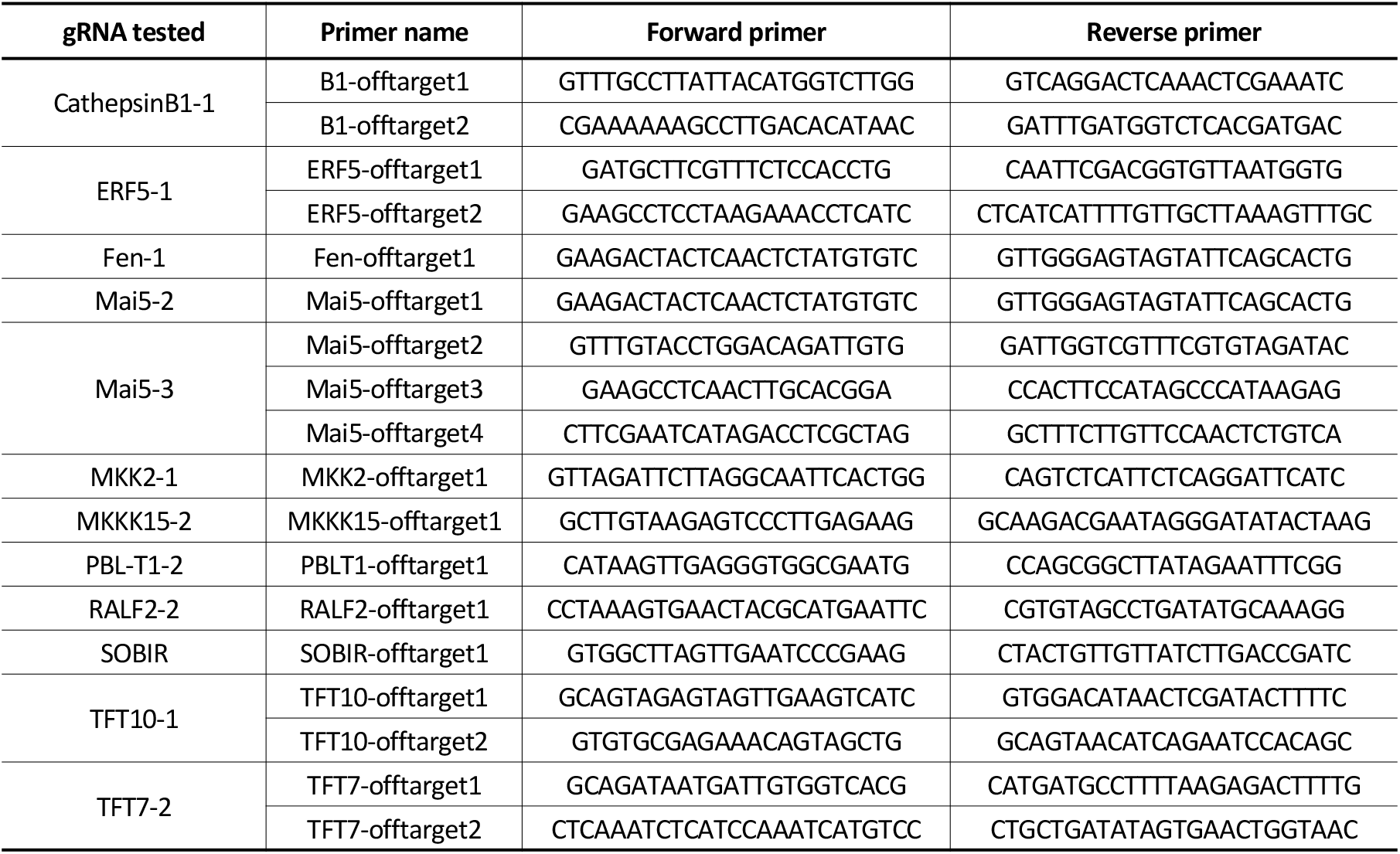
Primers used for off-target analysis (related to Table 3).

